# Classification of skin transcriptome reveals two molecular subtypes in hidradenitis suppurativa

**DOI:** 10.1101/2025.02.03.636243

**Authors:** Jing Wang, Heath Guay, Dan Chang

## Abstract

**Background and Aims:** Hidradenitis suppurativa (HS) is an understudied chronic inflammatory skin disease that is characterized by painful bumps and abscesses. Adalimumab and secukinumab are the only two approved biologics for the treatment of HS. Despite these advances in the treatment of disease, there remains significant unmet medical need, with many patients only achieving moderate improvements in disease, or continuing to experience flares. This suggests that there may be distinct patient subsets with unique molecule drivers of disease. The aim of this study is to identify and characterize the molecular subtypes of HS patients to further understand its heterogeneity.

**Methods:** Six public datasets with a total of 100 skin lesional samples were integrated to identify molecular subtypes and build a 36-gene classifier, which was validated by three independent lesional skin datasets. Two out of six training datasets generated from patients treated with adalimumab were used to identify the relationship between subtypes and response status.

**Results:** Two molecular subtypes were identified from the training datasets, in which subtype S1 had a higher response rate of adalimumab than subtype S2 in the two adalimumab treatment datasets. Subtype S1 was characterized by three gene modules associated with keratinization, development, and metabolism, and six cell types related to sebocytes, smooth muscle cells, endothelial cells, schwann cells, basal cells, and proliferating cells. Subtype S2 was associated with three modules related to immune response, wound healing, keratinization, and cell cycle, and three cell types related to T lymphocytes, dendritic cells, and fibroblasts. The two subtypes were replicated in three additional independent datasets.

**Conclusions:** This study discovered and validated two HS subtypes with different molecular mechanisms and drug response, which may aid interpretation of heterogeneous molecular and clinical information in HS patients.

## Introduction

Hidradenitis suppurativa (HS) is a complex, chronic inflammatory skin disease characterized by recurrent, painful nodules and abscesses of apocrine gland-bearing skin^1^. HS affects approximately 0.1%-1% of the population globally^2^, with large variations across different countries^3^. HS usually develops after puberty, with an average onset in the second or third decade of life and with a female predominance. Due to its chronic nature and frequently occurring relapses, HS has a more significant impact on the patients’ quality of life than that shown for other skin diseases^4^.

Despite the increased burden caused by HS alterations, the pathogenesis of HS remains incompletely understood. A key triggering factor is the occlusion of the hair follicle and cyst development^5^. The ruptured cyst will lead to an acute immune response and inflammation. Several studies hypothesized that the disease is also triggered by genetic (up to 38% of HS patients with a family history of the disease)^6^ and environmental factors (e.g. obesity and smoking)^7^. In addition, HS can occur with several co-morbid auto-immune diseases, particularly inflammatory bowel disease (IBD)^8^.

Studies implicating dysregulated immune responses in HS (e.g. elevations in tumor necrosis factor (TNF)-α, interleukin (IL)-1α/β, and IL-17) provide a rationale for therapies with biologics targeting these cytokines^9,10^. The TNF-α antagonist adalimumab is the first US FDA-approved therapy for this disease. IL17A antagonist secukinumab was also approved to treatment moderate to severe HS patients in 2023^11^. IL1α/β antagonist lutikizumab shows positive results in a HS phase 2 trial^12^. Currently, adalimumab and secukinumab demonstrate therapeutic response in up to 60% of patients^11,13^ and have the high numbers of flares among responders^14^, which suggests that there may be multiple distinct molecular drivers of this disease across patients. However, little has been done on the characterization of HS molecular subtypes. To our knowledge, only Navrazhina et al.^15^ identified two molecular subtypes from the skin transcriptomic dataset with 22 HS patients based on the expression of neutrophil associated gene lipocalin-2 (LCN2). They also found that genes up-regulated in LCN2-high subtype were enriched with inflammatory pathways (e.g. neutrophil chemotaxis and degranulation and IL-17-related pathways). It is still unknown whether this one-gene classification identified from a small sample size is reproducible in other datasets and whether LCN2 subtypes are associated with response status.

In this study, through integration of six HS transcriptomic datasets of a total of 100 skin lesional samples, we built the most comprehensive training cohort to identify the molecular subtypes of HS and validated them in three independent datasets. We assessed the association between subtypes and response status of adalimumab, and characterized the molecular and cellular mechanisms of each subtype.

## Materials and Methods

### Training and validation datasets

To build the training datasets, we searched Gene Expression Omnibus (GEO, https://www.ncbi.nlm.nih.gov/geo/) for transcriptomic studies with HS patients. After removing the studies with less than five lesional skin samples, six RNA sequencing (RNASeq) datasets (GSE151243^16^, GSE154773^17^, GSE155176^18^, GSE189266^19^, GSE213761^20^, and GSE249027^21^) were selected as the training datasets (Figure 1, Table 1, and supplementary Table S1). Only lesional samples from baseline were included in the training dataset. For each RNASeq dataset, the raw count matrix was downloaded from GEO. Three datasets (GSE72702^22^, GSE148027^23^, and GSE128637^24^) were selected as the validation datasets and the processed datasets were downloaded from GEO. Probe ID were mapped to the HUGO gene symbols based on a mapping table from GEO.

**Table 1.**
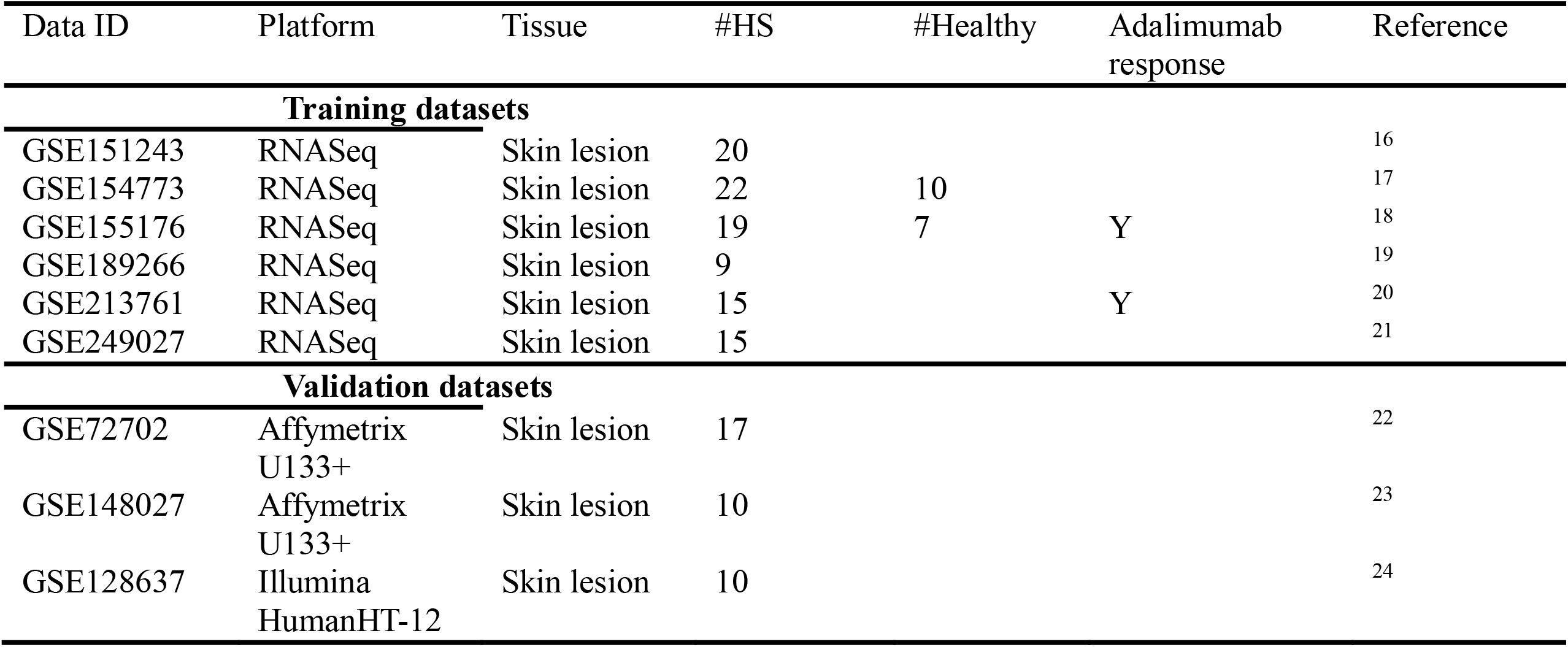
Training and validation datasets used in this study.

**Figure 1.**
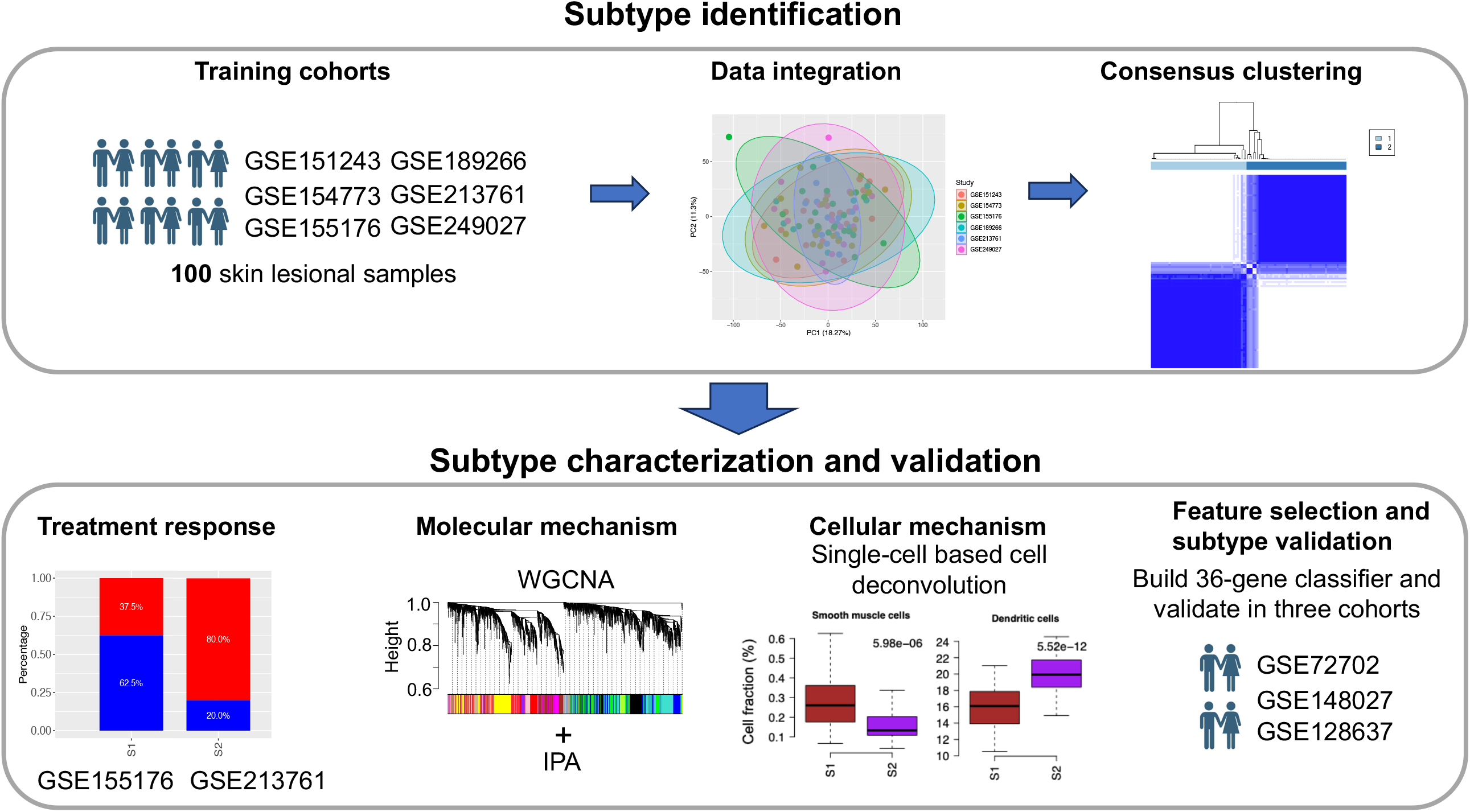
Overview of the analytical pipeline in this study. We integrated six HS datasets with 100 skin lesion samples to identify the subtypes by unsupervised consensus clustering method. Two datasets were used to evaluate the relationship between subtypes and adalimumab response. Then, WGCNA and cell deconvolution methods were used for revealing molecular and cellular mechanisms of subtypes. Subtype classifier was built and applied to three independent datasets for the validation.

### Training Data Integration and Subtype Identification

Based on 15,842 common genes among six training datasets, ComBat-seq^25^ was used to remove study-level batch effect and integrate six count matrices (supplementary Figure S1). After filtering the quantifiable genes by CPM (counts per million)>1 in at least 20% of samples, the integrated count matrix with 14,072 genes was normalized based on the TMM (trimmed mean of M-values) method^26^ and transformed by logarithmic function.

1,586 genes with MAD (median absolute deviation)>1 were used to identify the subtypes with R package ConsesusClusterPlus (version 1.62.0)^27^. Before clustering, the normalized training data matrix was scaled at gene level. Consensus clustering was performed using PAM (partition around medoids) method and a distance metric equals to 1 minus the Pearson’s correlation coefficient. A total of 1,000 permutation tests were performed with a subsampling ratio of 0.8. The maximum cluster number was set to 5. Then, four scores were used to define the optimal number of subtypes: proportion of ambiguous clustering^28^, Davies-Bouldin’s index^29^, Calinski-Harabasz index^30^, and average silhouette width^31^. The optimal number of subtypes had lower proportion of ambiguous clustering and Davies-Bouldin’s index and higher Calinski-Harabasz index and average silhouette width. Core samples of each subtype were defined as those with higher similarity to their own subtype than to any other subtypes and identified by silhouette width≥0.01^32^. The differential expressed genes (DEGs) between two subtypes were identified under |log2(Fold change)|>log2(1.5) and FDR<0.05 by R package limma (version 3.54.2)^33^.

### WGCNA Module Identification

Based on the DEGs between subtypes, WGCNA (weighted gene correlation network analysis) method^34^ was used to identify co-expressed gene modules in the integrated and normalized training dataset. Scale-free topology fit index (R^^2^) for each soft thresholding power was calculated by the pickSoftThreshold function in R package WGCNA (version 1.72-1). The power 4 with R^^2^=0.916 was selected to calculate the adjacency matrix (supplementary Figure S2). Because genes with a negative correlation did not correspond to the functional similarity^35^, only positive correlations were included in the adjacency matrix. Topology overlap matrix (TOM)^34^ was then calculated based on the adjacency matrix. Finally, the modules were identified based on TOM using the cutreeDyanmic function in WGCNA package with minClusterSize=100. Module expression was computed using the GSVA (gene set variation analysis) method^36^. Differential module expression between subtypes was calculated by Wilcoxon rank-sum test and p-value was adjusted by the Benjamini-Hochberge method. Significant modules were identified under FDR<0.05. Pathway analysis for each module was performed using Ingenuity Pathway Analysis (IPA, www.ingenuity.com). The significant canonical pathways were identified under FDR<0.05 and the number of overlapping genes between module genes and genes in the IPA pathway≥5. Upstream regulators of each module were predicted by IPA under FDR<0.05.

### Deep-Learning Based Cell Fraction Estimation Using Single-Cell RNA Sequencing Data

HS single-cell RNA sequencing (scRNA-seq) dataset GSE154775 had the largest number of HS samples compared with other HS scRNA-seq datasets, which was used to estimate the cell fractions of bulk transcriptomic samples. For each scRNA-seq dataset, cell ranger outputs were downloaded from GEO and processed by harmony (version 1.2.1)^37^ and Seurat (version 4.3.0)^38^ R packages. Cell type annotation was performed by the GPTCelltype (version 1.0.1) R package. Then, the Scaden (single cell-assisted deconvolutional deep neural network) method developed by Menden et al.^39^ was used to deconvolute the normalized training dataset based on 12 cell types from GSE154775. A p-value comparing cell fractions between subtypes was calculated by Wilcoxon rank-sum test and adjusted by the Benjamini-Hochberge method. Significant cell types were identified under the mean of cell fractions in at least one subtype>0.1%, and FDR<0.05.

### Construction of Subtype Classifier

The top 200 DEGs with minimal p-values were used to perform the feature selection by the recursive feature elimination (RFE) method with a random forest model [24]. Based on number of genes from 1 to 200, RFE function built the random forest model with three-time repeated five-fold cross-validation to calculate the AUROC (the area under the ROC curve). Because AUROC was stable after 36 genes (supplementary Figure S3), top 36 genes with the largest overall scores (supplementary Table S2) were selected to build the final random forest model. The subtype of each sample in the validation datasets was predicted by selecting the subtype with the largest probability returned from the final model.

### Data Availability

GEO bulk and single-cell datasets can be downloaded from public data portals.

## Results

### Two Molecular Subtypes in HS

Two clusters identified from consensus clustering method had the lowest proportion of ambiguous clustering score, lowest Davies-Bouldin index, highest Calinski-Harabasz index and highest average silhouette width (Figure 2A) compared with other cluster numbers. These consistent statistics indicated that the 100 HS skin lesional samples in the training dataset could be robustly divided into two subtypes. Then, 5 samples were discarded due to lower similarity within one subtype than the other (Figure 2B). Principal component analysis (PCA) of the 95 core samples revealed significant difference between two subtypes (p-value<2.2E-16, Figure 2C) and the distribution of two subtypes across six studies included in the training dataset were balanced (p-value=0.73, supplementary Figure S4). To test the confounding factors for the subtype identification, we checked whether the subtypes were associated with age, BMI (body mass index) and smoking status in two studies GSE151243 and GSE213761. As shown in supplementary Figure S5, there was no relationship between subtypes and age in both datasets. However, there was inconsistent results for BMI and smoking status. Subtype S2 in GSE213761 but not GSE151243 had significantly higher BMI while subtype S2 in GSE151243 but not GSE213761 was represented by more non-smokers. GSE213761 also included Hurley stage information, and we found that although there was no significant relationship between subtypes and different stages (p=0.29, supplementary Figure S5B), subtype S1 included 75% Stage II patients and 25% Stage III patients compared with 40% and 60% in subtype S2. Thus, subtype S1 may include more moderate HS patients. Two studies GSE155176 and GSE213761 included the drug response information for adalimumab, which were used to assess the association between subtypes and response status. In GSE213761, we found that subtype S1 included 62.5% responders defined by HiSCR (hidradenitis suppurativa clinical response)^40^ while this percentage decreased to 20% in subtype S2 (Figure 2D). In GSE155176, the percentage of responders defined by HiSCR in both subtypes were 66.7% and 36.4%, respectively (Figure 2E). Although the p-values were not significant due to the small sample size, these consistent results still supported that subtype S1 enriched with responders of adalimumab while subtype S2 was associated with non-responders.

**Figure 2.**
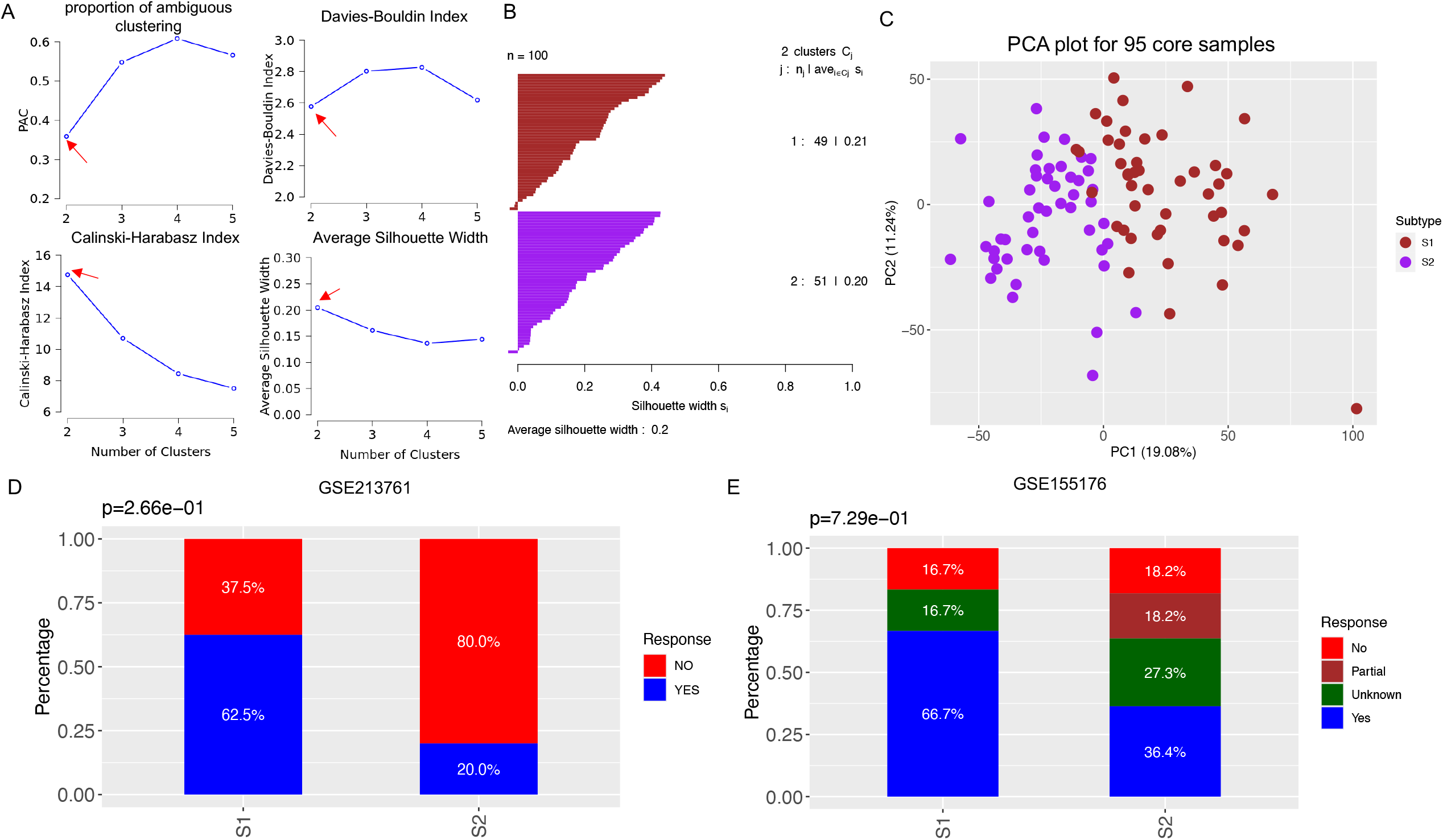
Identification of HS subtypes. (A) Relationship between number of clusters and four scores. The red arrow in each figure represents the optimal cluster number. (B) Silhouette plot for removing the outlier samples. (C) PCA plot for 95 core samples. (D-E) Relationship between subtypes and adalimumab response. The p value was calculated by the Fisher’s exact test.

Navrazhina et al. ^15^ identified two HS subtypes based on LCN2 expression and LCN2-high subtype was characterized as highly inflammatory HS subtype. Comparing with the subtypes in this study, we found that 70.2% LCN2-high subtype were included in the subtype S2 while 66.7% LCN2-low subtype belonged to subtype S1 (supplementary Figure S6A). In GSE213761, LCN2-high and -low subtypes had 50.0% and 44.4% response rate, respectively (supplementary Figure S6B). In contrast, LCN2-high subtype had lower response rate in GSE155176 (40.0%) than LCN2-low subtype (57.1%) (supplementary Figure S6C). Thus, there was no relationship between LCN2 subtypes and adalimumab response in the cohorts studied here.

### Molecular Mechanisms of the Two Subtypes

Comparing subtype S1 with subtype S2 samples, 1,046 up-regulated genes and 1,220 down-regulated genes were identified under |log2(Fold change)|>log2(1.5) and FDR<0.05 (supplementary Figure S7 and Table S3). Six WGCNA modules (modules 1-6, supplementary Table S4) were identified based on the DEGs, in which three modules were associated with subtype S1 and three modules were related to subtype S2 (FDR<0.05, Figure 3A, supplementary Table S5).

**Figure 3.**
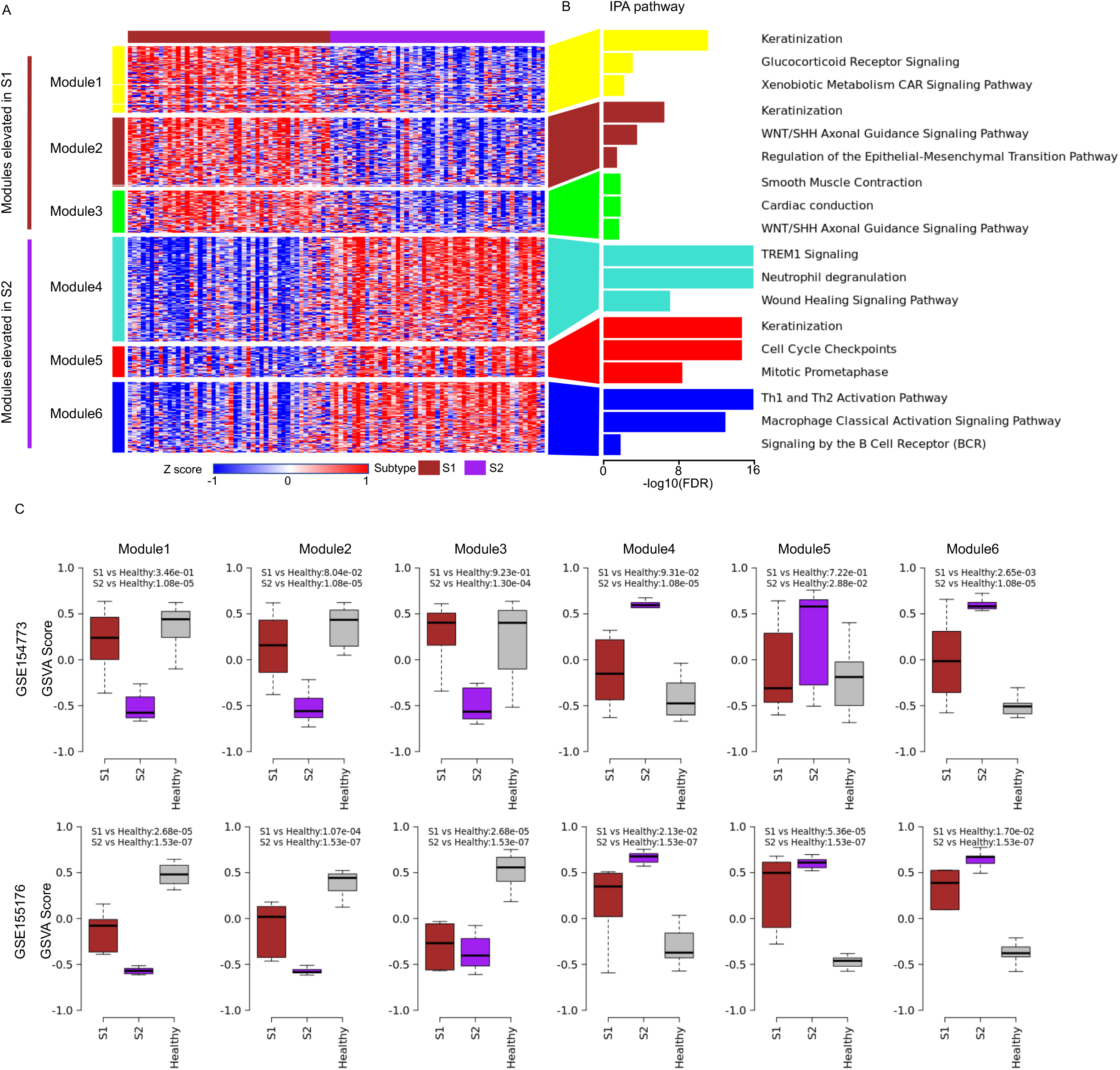
Molecular mechanisms of HS subtypes. (A) Heatmap of z-transformed expression of genes from each of six modules in the integrated training dataset. The samples were ordered by the subtypes. (B) Functional annotation of genes in each module based on IPA. Three representative enriched pathways were shown, sorted by p-value. (C) Comparison of GSVA scores of six modules among subtype S1, S2, and healthy samples in GSE154773 and GSE155176. The p-values were calculated based on Wilcoxon rank sum test.

Figure 3B showed the representative IPA pathways enriched in each module (all enriched IPA pathways can be found in supplementary Table S6). Because HS is closely linked with aberrant keratinization^41^, keratinization pathway was highly enriched in both subtypes (modules 1-2 for subtype S1 and module 5 for subtype S2). Meanwhile, subtype S1-related module 1 also reflected metabolism pathways (e.g. glucocorticoid receptor signaling pathway and xenoblotic metabolism CAR signaling pathway) while modules 2 and 3 were highly related to the cell signaling and development of skin tissue (e.g. WNT/SHH axonal guidance signaling pathway and smooth muscle contraction). Subtype S2-related module 4 was mainly associated with innate immune pathways (e.g. TREM1 signaling and neutrophil degranulation) while module 6 was related to adaptive immune response (e.g. Th1 and Th2 activation pathway, macrophage classical activation signaling pathway, and signaling by the B Cell Receptor). Module 5 was enriched with cell cycle pathways (e.g. cell cycle checkpoints and mitotic prometaphase). We also found that wound healing signaling pathway was significant in modules 4-6 but not modules 1-3 (supplementary Table S5).

GSE154773 and GSE155176 included the healthy samples (Table 1), and we compared the GSVA scores of six modules among subtype S1, S2, and healthy samples. As shown in Figure 3C, the difference between subtype S1 and healthy samples were smaller than those between subtype S2 and healthy samples for six modules, which also implied that subtype S1 may include more moderate patients.

Up-stream regulator analysis identified 14 regulators for at least two S1 modules and 262 regulators related to at least two S2 modules (Figure 4 and supplementary Table S7). For example, androgen receptor (AR) was the up-stream regulator for S1-related modules 2 and 3, which played important role in HS pathogenesis^42^.

**Figure 4.**
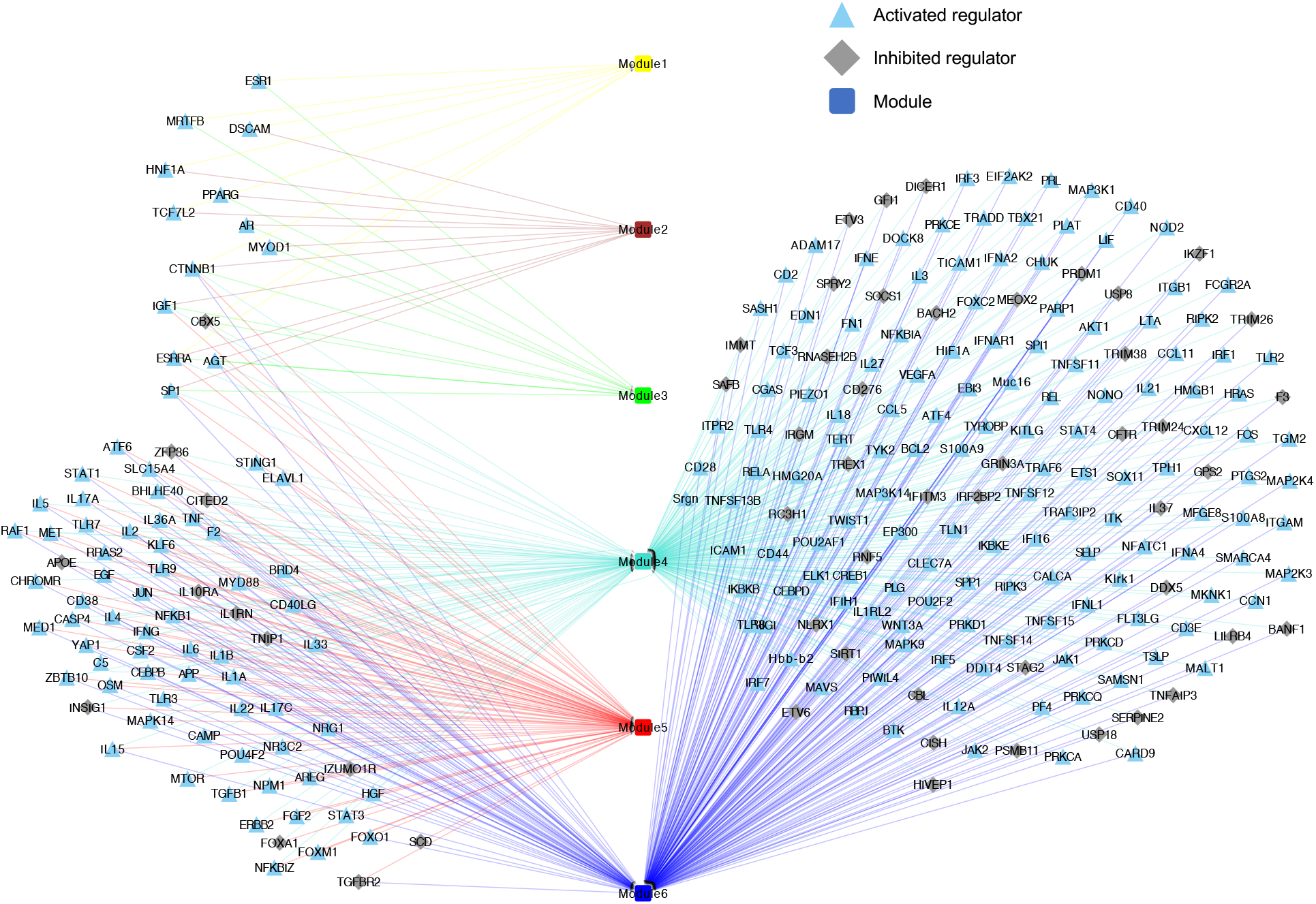
Up-stream regulators of six modules. Triangle, diamond, and square represent activated regulator, inhibited regulator, and module, respectively. Regulators related to at least 2 modules were visualized in the figure.

### Cellular Mechanisms of the Two Subtypes

As shown in Figure 5, cell deconvolution analysis using the scRNA-seq dataset GSE154775 identified six cell types that had significantly higher cell fractions in subtype S1 compared with subtype S2 (FDR<0.05). The six cell types included sebocytes, smooth muscle cells, endothelial cells, schwann cells, basal cells, and proliferating cells. Two immune cells (dendritic cells and T lymphocytes) and fibroblasts had higher cell fractions in subtype S2. This result provided the consistent conclusion between molecular and cellular mechanisms of the two subtypes. Furthermore, keratinocytes had over 20% cell fractions in most of samples and had no difference between two subtypes, which was also consistent with the molecular mechanisms.

**Figure 5.**
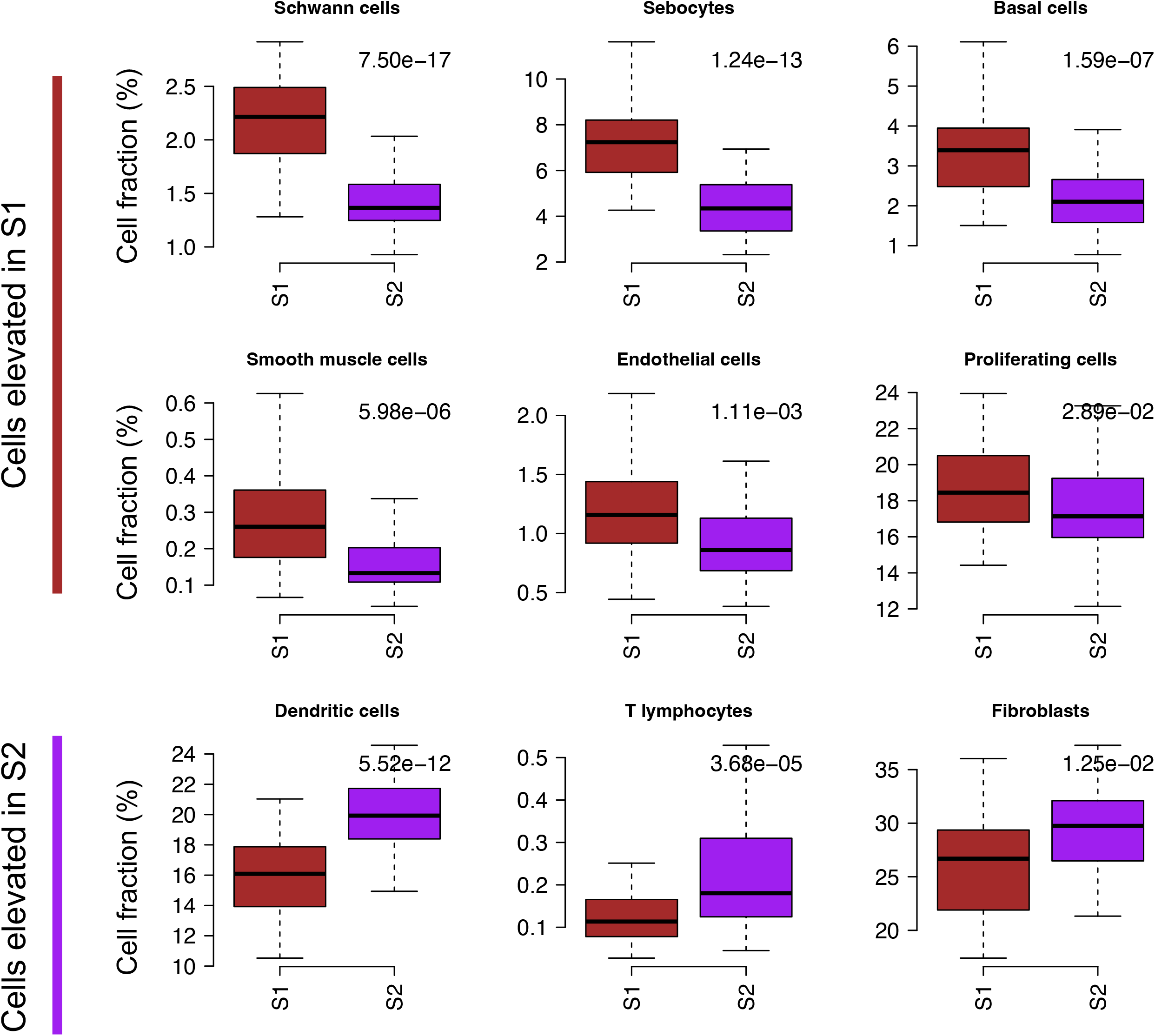
Cellular mechanisms of HS subtypes. Cell types elevated in subtype S1 and S2. The p-value was calculated based on Wilcoxon rank sum test.

### Validation of Two Subtypes

To validate our two-subtype classification, a classifier with AUROC>0.99 was built based on 36 genes selected from RFE feature selection method (supplementary Figure S3 and Table S2). Among 36 genes, 7 genes were from the metabolism module (module1), 7 genes were from skin development module (modules 2-3), and 22 genes were from immune and wound healing modules (modules 4 and 6).

Applying our 36-gene classifier to three independent validation datasets, the two subtypes can be robustly rediscovered in these datasets (Figure 6A-C). We also performed consensus clustering for GSE72702 and the clusters were consistent with predicted ones (supplementary Figure S8).

**Figure 6.**
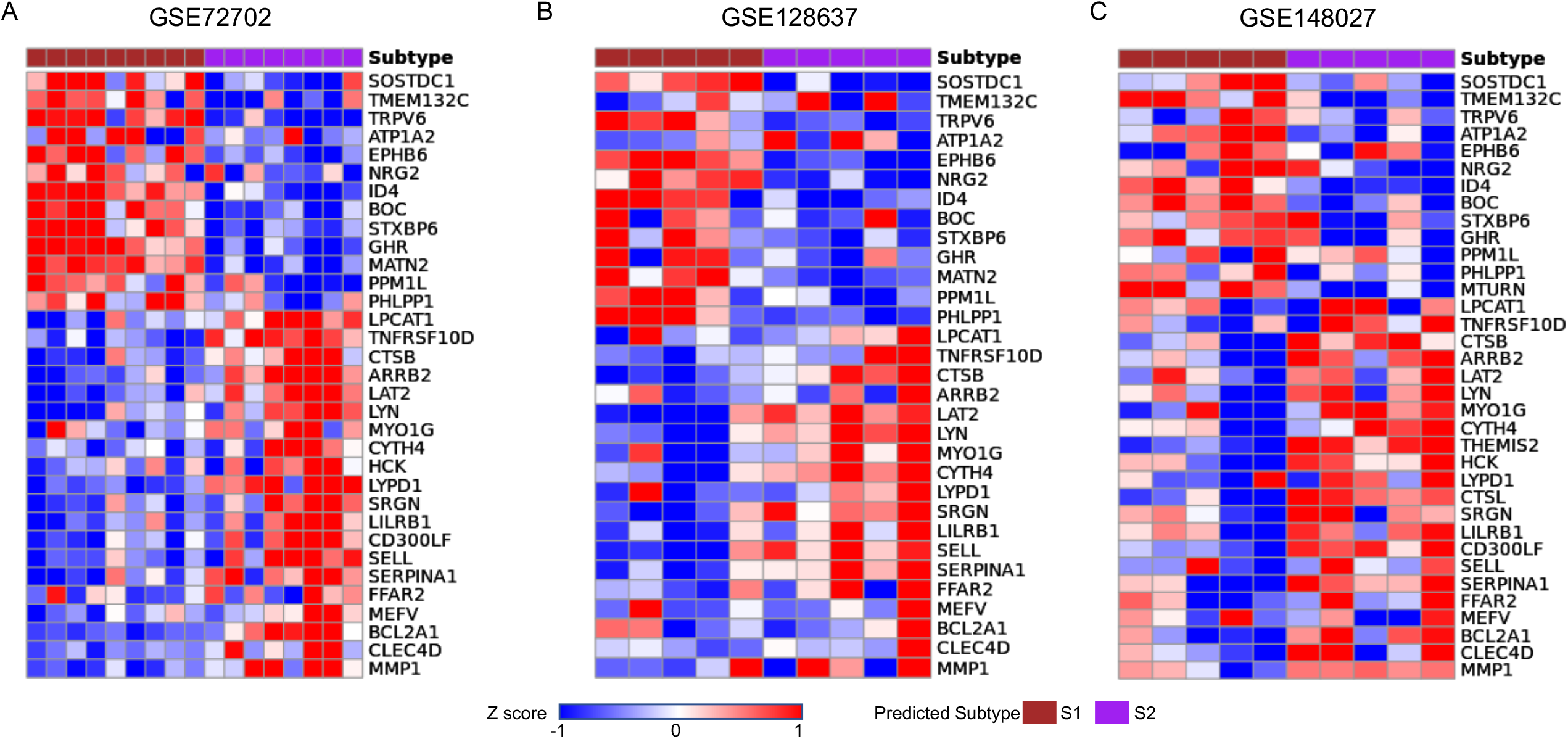
Subtype validation. (A-C) Heatmap of z-transformed expression of 36 genes in three independent datasets. The 36 genes were ordered by their log2(Fold change) between subtype S1 and S2 in the integrated training dataset and the samples were ordered by the predicted subtypes.

## Discussion

As the only two US FDA-approved therapies for HS, adalimumab and secukinumab show a response in up to 60% of patients. There is still high unmet medical need of these patients, and the heterogeneous response to existing therapies suggests that there may be multiple distinct molecular drivers of disease across unique subsets of patients. Based on the most comprehensive training datasets with 100 HS skin lesional samples, we identified two molecular subtypes, which have been validated by three independent datasets using a 36-gene classifier we developed. We also found that subtype S1 enriched with adalimumab responders while subtype S2 was associated with non-responders.

Only one study has proposed molecular subtypes for HS based on transcriptomic dataset^15^. Based on the neutrophil associated gene LCN2, Navrazhina et al. identified LCN2-high and LCN2-low subtypes and found that genes up-regulated in LCN2-high subtype were enriched with inflammatory pathways (e.g. neutrophil chemotaxis and degranulation and IL-17-related pathways). Comparing LCN2 subtypes with the subtypes in this study, we found that LCN2-high subtype was associated with subtype S2 while LCN2-low subtype belonged to subtype S1 (supplementary Figure S6A). We also compared the expression of neutrophil genes (LCN2, CXCL1, CXCL8, CD177, and CSF3) between subtype S1 and S2 and found that all these genes had significantly higher expression in subtype S2 than S1 (supplementary Table S3). Thus, our subtypes were partially in agreement with LCN2 subtypes. However, when comparing with response status of adalimumab, we found that subtype S1 had higher response rate than subtype S2 in GSE213761 and GSE155176 while there was no relationship between LCN2 subtypes and response status in two datasets (supplementary Figure S6B-C). Thus, our subtypes, identified from larger training datasets of bulk transcriptomics, represent significant improvement in understanding the molecular subtypes of HS.

Lowe et al.^18^ compared adalimumab responders with non-responders based on GSE155176 baseline samples and found that genes significantly increased in responders were related to epithelial and skin development while genes elevated in non-responders involved immune related pathways. Thus, they indicated that HS patients with more heightened global inflammation at baseline were less likely to respond to adalimumab. In this study, we found that subtype S1 had higher adalimumab response rate than subtype S2. Because subtype S1 modules were highly related to the metabolism and skin development while subtype S2 modules involved immune, wound healing, and cell cycle pathways, our findings were consistent with Lowe et al.. Meanwhile, Lowe et al.^18^ and Hambly et al.^20^ also found that a heightened B cell infiltrate in skin at baseline may be less likely to respond to adalimumab. Because scRNA-seq dataset used for the cell deconvolution only included 368 plasma cells, cell fractions of plasma cells predicted from deep learning model were very low for 95 training samples. However, subtype S2 related module 4 and 6 were enriched with B cell related pathways (supplementary Table S6) and CD19, CD38, and CXCR5 had higher expression in subtype S2 than S1 (FDR=4.3E-05 and 1.3E-04, supplementary Table S2), which was also consistent with Lowe et al. findings.

Based on meta-analysis from 93,601 unique participants from multiple observational studies, Chen et al.^8^ examined the risk of IBD in patients with HS and found the significant comorbidity association between HS and IBD. However, it is still unknown whether HS and IBD share the similar mechanisms or subtypes. Recently, we identified two subtypes shared by Crohn’s disease and ulcerative colitis: immune & tissue remodeling subtype and metabolism subtype^43^, which has similar molecular profiles as the two HS subtypes discovered here. To test the similarity between HS and IBD subtypes, we applied our HS 36-gene classifier discovered here to the 827 IBD colon training samples used in ^43^. As shown in supplementary Figure S9, HS subtype S1 was significantly enriched with IBD metabolism subtype while HS subtype S2 was highly associated with IBD immune & tissue remodeling subtype. Interestingly, in addition to the similarity in molecular mechanisms, the IBD metabolism subtype is enriched with responders to anti-TNF therapy while IBD immune & tissue remodeling subtype was associated with non-responders, which is also consistent with the response to HS subtypes to adalimumab discovered here. These results implied very similar molecular mechanism between IBD and HS, a novel discovery that should be further investigated in the future studies.

Because HS can be triggered by the environmental factors (e.g. obesity and smoking)^7^, we checked whether BMI and smoking status were cofounding factors of the subtype identification and did not find the consistent results (see supplementary Figure S6). The subtype S1 related module 1 was associated with glucocorticoid receptor signaling, which may imply that the patients in subtype S1 was treated by glucocorticoid before sample collection. However, because of the limited meta data provided by the training datasets, we can not test the associations.

Genetic factors clearly play an important role in causing HS. Sun et al.^44^ performed GWAS analysis based on 720 HS patients and found two lead variants near the SOX9 and KLF5. However, SOX9 and KLF5 were not DEGs between subtype S1 and S2 or module up-stream regulators (supplementary Table S2 and S6). Moltrasio et al.^45^ also collected 16 genes that had genetic changes involved in all forms of HS. Among these 16 genes, five genes (PSTPIP1, MEFV, NLRP3, NLRC4, GJB2, and KRT6) had significantly higher expression in subtype S2, but no gene was associated with subtype S1, implying that these genetic factors may be more associated with the inflammation mechanisms of the subtype S2 patients.

## Conclusion

In this study, based on 100 skin lesion samples, we identified two robust HS subtypes. Subtype S1 was associated with keratinization, development, and metabolism while subtype S2 was associated with immune response, wound healing, keratinization, and cell cycle. Furthermore, subtype S1 enriched with adalimumab responders while subtype S2 was associated with non-responders. These findings can aid interpretation of molecular heterogeneity in HS patients.

## Reference

1. Sabat R, Jemec GBE, Matusiak L, et al. Hidradenitis suppurativa. Nat Rev Dis Primers 2020;6:18.

2. Putkowski S. National organization for rare disorders (nord): Providing advocacy for people with rare disorders. NASN Sch Nurse 2010;25:38–41.

3. Jfri A, Nassim D, O’Brien E, et al. Prevalence of hidradenitis suppurativa: A systematic review and meta-regression analysis. JAMA Dermatol 2021;157:924–31.

4. von der Werth JM, Jemec GB. Morbidity in patients with hidradenitis suppurativa. Br J Dermatol 2001;144:809–13.

5. von Laffert M, Helmbold P, Wohlrab J, et al. Hidradenitis suppurativa (acne inversa): Early inflammatory events at terminal follicles and at interfollicular epidermis. Exp Dermatol 2010;19:533–7.

6. van Straalen KR, Prens EP, Willemsen G, Boomsma DI, van der Zee HH. Contribution of genetics to the susceptibility to hidradenitis suppurativa in a large, cross-sectional dutch twin cohort. JAMA Dermatol 2020;156:1359–62.

7. Sartorius K, Emtestam L, Jemec GB, Lapins J. Objective scoring of hidradenitis suppurativa reflecting the role of tobacco smoking and obesity. Br J Dermatol 2009;161:831–9.

8. Chen WT, Chi CC. Association of hidradenitis suppurativa with inflammatory bowel disease: A systematic review and meta-analysis. JAMA Dermatol 2019;155:1022–7.

9. van der Zee HH, de Ruiter L, van den Broecke DG, et al. Elevated levels of tumour necrosis factor (tnf)-alpha, interleukin (il)-1beta and il-10 in hidradenitis suppurativa skin: A rationale for targeting tnf-alpha and il-1beta. Br J Dermatol 2011;164:1292–8.

10. Kelly G, Hughes R, McGarry T, et al. Dysregulated cytokine expression in lesional and nonlesional skin in hidradenitis suppurativa. Br J Dermatol 2015;173:1431–9.

11. https://www.cosentyx.com/hidradenitis-suppurativa/index.

12. AbbVie. Lutikizumab showed positive results in a phase 2 trial of adults with moderate to severe hidradenitis suppurativa as program advances to phase 3. https://news.abbvie.com/2024-01-08-Lutikizumab-Showed-Positive-Results-in-a-Phase-2-Trial-of-Adults-with-Moderate-to-Severe-Hidradenitis-Suppurativa-as-Program-Advances-to-Phase-3, 2024.

13. Aarts P, van Huijstee JC, van der Zee HH, et al. Adalimumab in conjunction with surgery compared with adalimumab monotherapy for hidradenitis suppurativa: A randomized controlled trial in a real-world setting. J Am Acad Dermatol 2023;89:677–84.

14. Caposiena Caro RD, Cannizzaro MV, Tartaglia C, Bianchi L. Clinical response rate and flares of hidradenitis suppurativa in the treatment with adalimumab. Clin Exp Dermatol 2020;45:438–44.

15. Navrazhina K, Garcet S, Zheng X, et al. High inflammation in hidradenitis suppurativa extends to perilesional skin and can be subdivided by lipocalin-2 expression. J Allergy Clin Immunol 2022;149:135–44 e12.

16. Freudenberg JM, Liu Z, Singh J, et al. A hidradenitis suppurativa molecular disease signature derived from patient samples by high-throughput rna sequencing and re-analysis of previously reported transcriptomic data sets. PLoS One 2023;18:e0284047.

17. Gudjonsson JE, Tsoi LC, Ma F, et al. Contribution of plasma cells and b cells to hidradenitis suppurativa pathogenesis. JCI Insight 2020;5.

18. Lowe MM, Naik HB, Clancy S, et al. Immunopathogenesis of hidradenitis suppurativa and response to anti-tnf-alpha therapy. JCI Insight 2020;5.

19. Navrazhina K, Frew JW, Grand D, et al. Interleukin-17ra blockade by brodalumab decreases inflammatory pathways in hidradenitis suppurativa skin and serum. Br J Dermatol 2022;187:223–33.

20. Hambly R, Gatault S, Smith CM, et al. B-cell and complement signature in severe hidradenitis suppurativa that does not respond to adalimumab. Br J Dermatol 2023;188:52–63.

21. Ben Abdallah H, Bregnhoj A, Iversen L, Johansen C. Transcriptomic analysis of hidradenitis suppurativa: A unique molecular signature with broad immune activation. Int J Mol Sci 2023;24.

22. Blok JL, Li K, Brodmerkel C, Jonkman MF, Horvath B. Gene expression profiling of skin and blood in hidradenitis suppurativa. Br J Dermatol 2016;174:1392–4.

23. Penno CA, Jager P, Laguerre C, et al. Lipidomics profiling of hidradenitis suppurativa skin lesions reveals lipoxygenase pathway dysregulation and accumulation of proinflammatory leukotriene b4. J Invest Dermatol 2020;140:2421–32 e10.

24. Shanmugam VK, Jones D, McNish S, Bendall ML, Crandall KA. Transcriptome patterns in hidradenitis suppurativa: Support for the role of antimicrobial peptides and interferon pathways in disease pathogenesis. Clin Exp Dermatol 2019;44:882–92.

25. Zhang Y, Parmigiani G, Johnson WE. Combat-seq: Batch effect adjustment for rna-seq count data. NAR Genom Bioinform 2020;2:lqaa078.

26. Robinson MD, Oshlack A. A scaling normalization method for differential expression analysis of rna-seq data. Genome Biol 2010;11:R25.

27. Wilkerson MD, Hayes DN. Consensusclusterplus: A class discovery tool with confidence assessments and item tracking. Bioinformatics 2010;26:1572–3.

28. Senbabaoglu Y, Michailidis G, Li JZ. Critical limitations of consensus clustering in class discovery. Sci Rep 2014;4:6207.

29. Davis DL, Bouldin DW. A cluster separation measure. IEEE Transactions on Pattern Analysis and Machine Intelligence 1979;2:224–7.

30. Calinski T, Harabasz J. A dendrite method for cluster analysis. Communications in Statistics 1974;3:1–27.

31. Rousseeuw PJ. Silhouettes: A graphical aid to the interpretation and validation of cluster analysis. Journal of Computational and Applied Mathematics 1987;20:53–65.

32. Verhaak RG, Hoadley KA, Purdom E, et al. Integrated genomic analysis identifies clinically relevant subtypes of glioblastoma characterized by abnormalities in pdgfra, idh1, egfr, and nf1. Cancer Cell 2010;17:98–110.

33. Ritchie ME, Phipson B, Wu D, et al. Limma powers differential expression analyses for rna-sequencing and microarray studies. Nucleic Acids Res 2015;43:e47.

34. Langfelder P, Horvath S. Wgcna: An r package for weighted correlation network analysis. BMC Bioinformatics 2008;9:559.

35. Wang J, Ma Z, Carr SA, et al. Proteome profiling outperforms transcriptome profiling for coexpression based gene function prediction. Mol Cell Proteomics 2017;16:121–34.

36. Hanzelmann S, Castelo R, Guinney J. Gsva: Gene set variation analysis for microarray and rna-seq data. BMC Bioinformatics 2013;14:7.

37. Korsunsky I, Millard N, Fan J, et al. Fast, sensitive and accurate integration of single-cell data with harmony. Nat Methods 2019;16:1289–96.

38. Hao Y, Hao S, Andersen-Nissen E, et al. Integrated analysis of multimodal single-cell data. Cell 2021;184:3573–87 e29.

39. Menden K, Marouf M, Oller S, et al. Deep learning-based cell composition analysis from tissue expression profiles. Sci Adv 2020;6:eaba2619.

40. Kimball AB, Sobell JM, Zouboulis CC, et al. Hiscr (hidradenitis suppurativa clinical response): A novel clinical endpoint to evaluate therapeutic outcomes in patients with hidradenitis suppurativa from the placebo-controlled portion of a phase 2 adalimumab study. J Eur Acad Dermatol Venereol 2016;30:989–94.

41. Nomura T. Hidradenitis suppurativa as a potential subtype of autoinflammatory keratinization disease. Front Immunol 2020;11:847.

42. Riis PT, Ring HC, Themstrup L, Jemec GB. The role of androgens and estrogens in hidradenitis suppurativa - a systematic review. Acta Dermatovenerol Croat 2016;24:239–49.

43. Wang J, Guay H, Chang D. Crohn’s disease and ulcerative colitis share two molecular subtypes with different mechanisms and drug response. J Crohns Colitis 2024.

44. Sun Q, Broadaway KA, Edmiston SN, et al. Genetic variants associated with hidradenitis suppurativa. JAMA Dermatol 2023;159:930–8.

45. Moltrasio C, Tricarico PM, Romagnuolo M, Marzano AV, Crovella S. Hidradenitis suppurativa: A perspective on genetic factors involved in the disease. Biomedicines 2022;10.

