## Supplementary Figures for "Classification of skin transcriptome reveals two molecular subtypes in hidradenitis suppurativa"

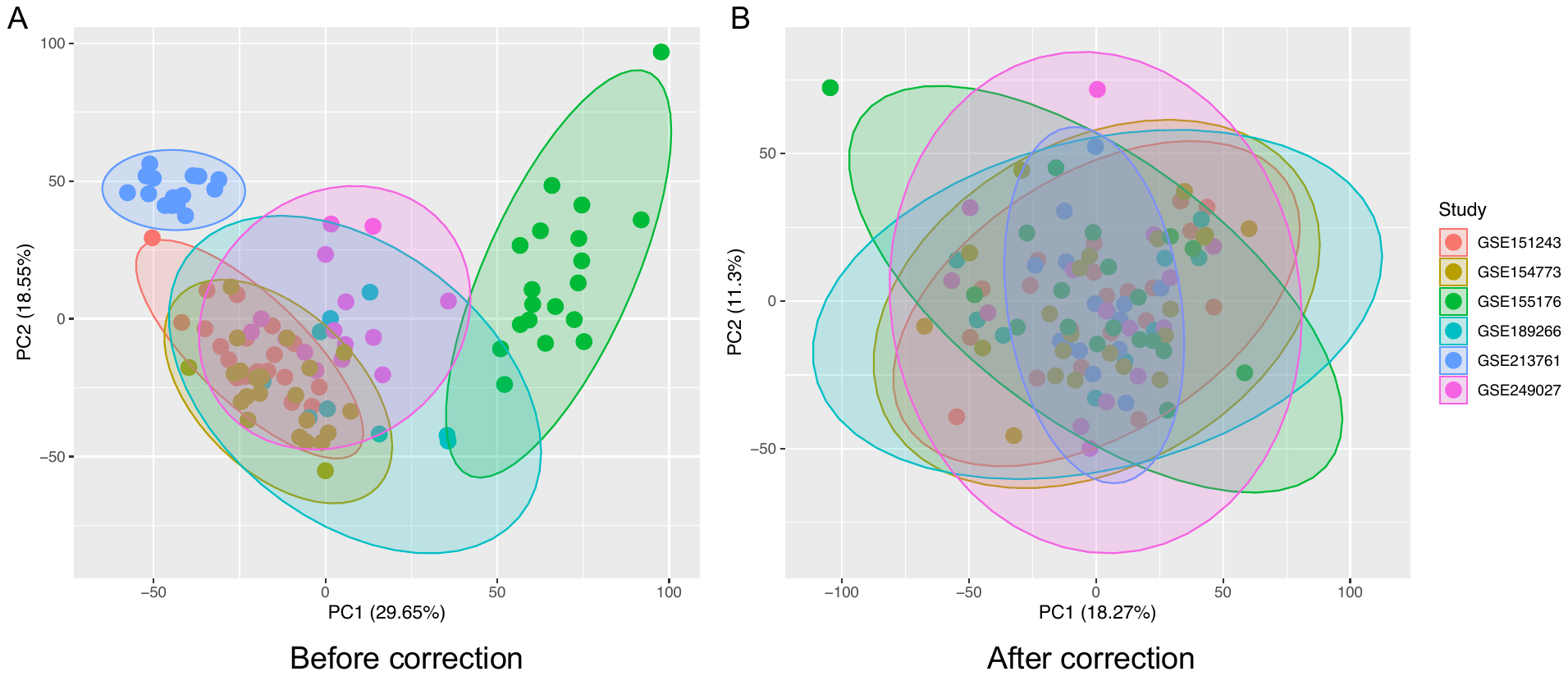

Supplementary Figure S1. PCA plots for six training dataset integration without (A) or with study-level batch correction (B). Colors represent different studies.

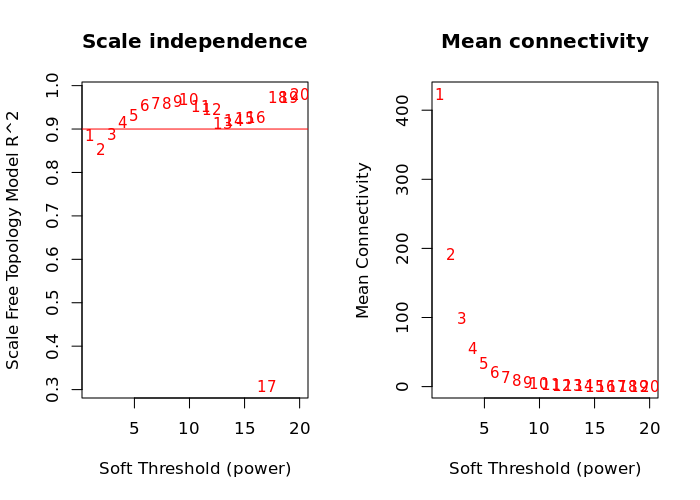

Supplementary Figure S2. Power selection for the WGCNA analysis based on scale-free topology fit index.

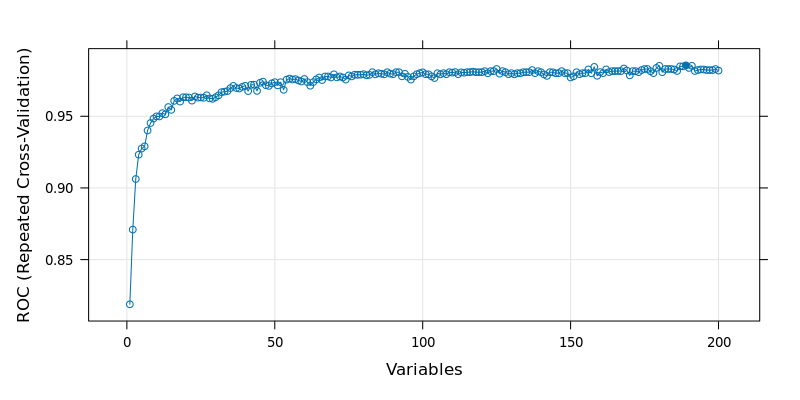

Supplementary Figure S3. Feature selection based on top200 DEGs. Because AUROC was stable after 36 genes, top 36 genes were selected to build a classifier for the subtype prediction.

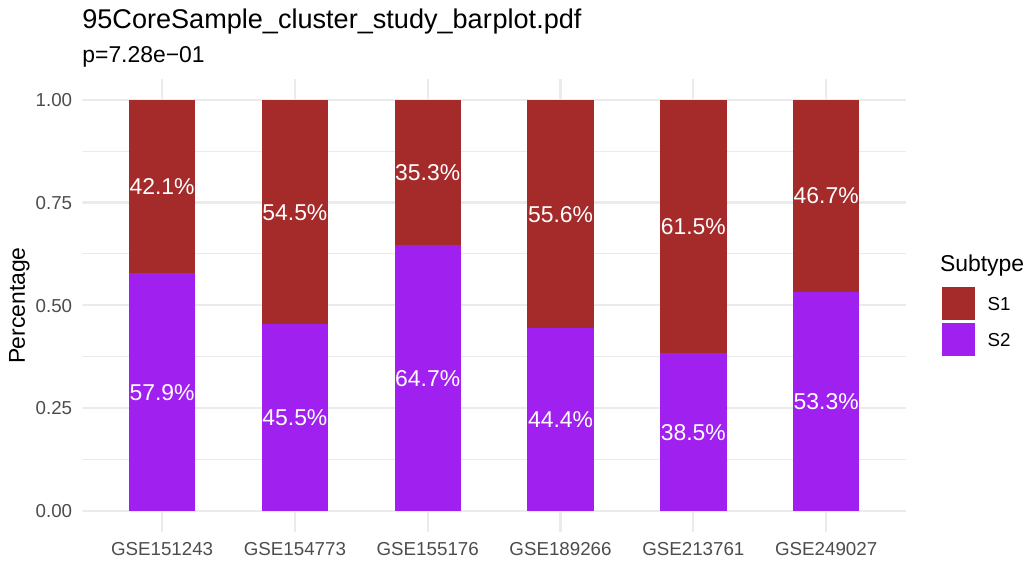

Supplementary Figure S4. Distribution of two subtypes across six training datasets. The p-value was calculated by the Fisher’s exact test.

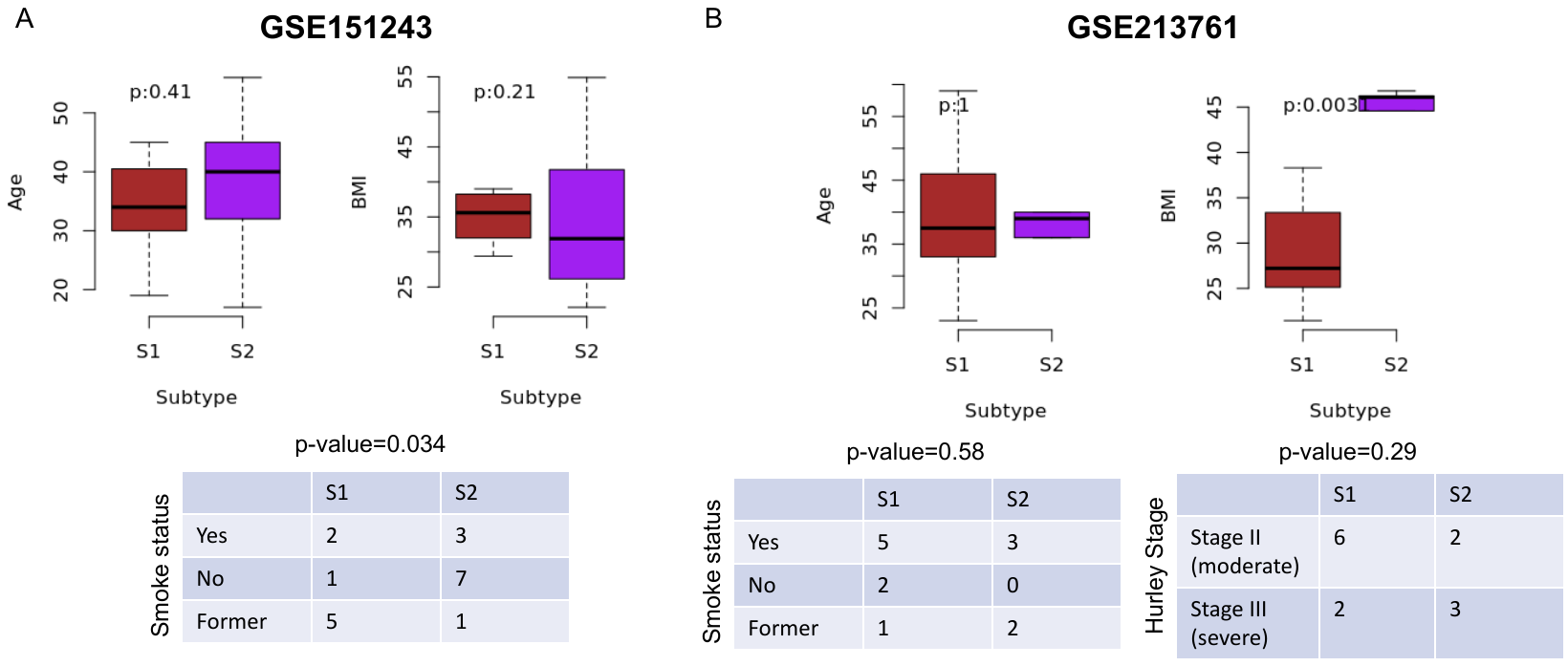

Supplementary Figure S5. The relationship between two subtypes and age/BMI/smoking status/Hurley Stage in GSE151243 (A) and GSE213761 (B). The p-values for age and BMI were calculated by Wilcoxon rank sum test. The p-values for smoking status and Hurley stage were calculated by Fisher’s exact test.

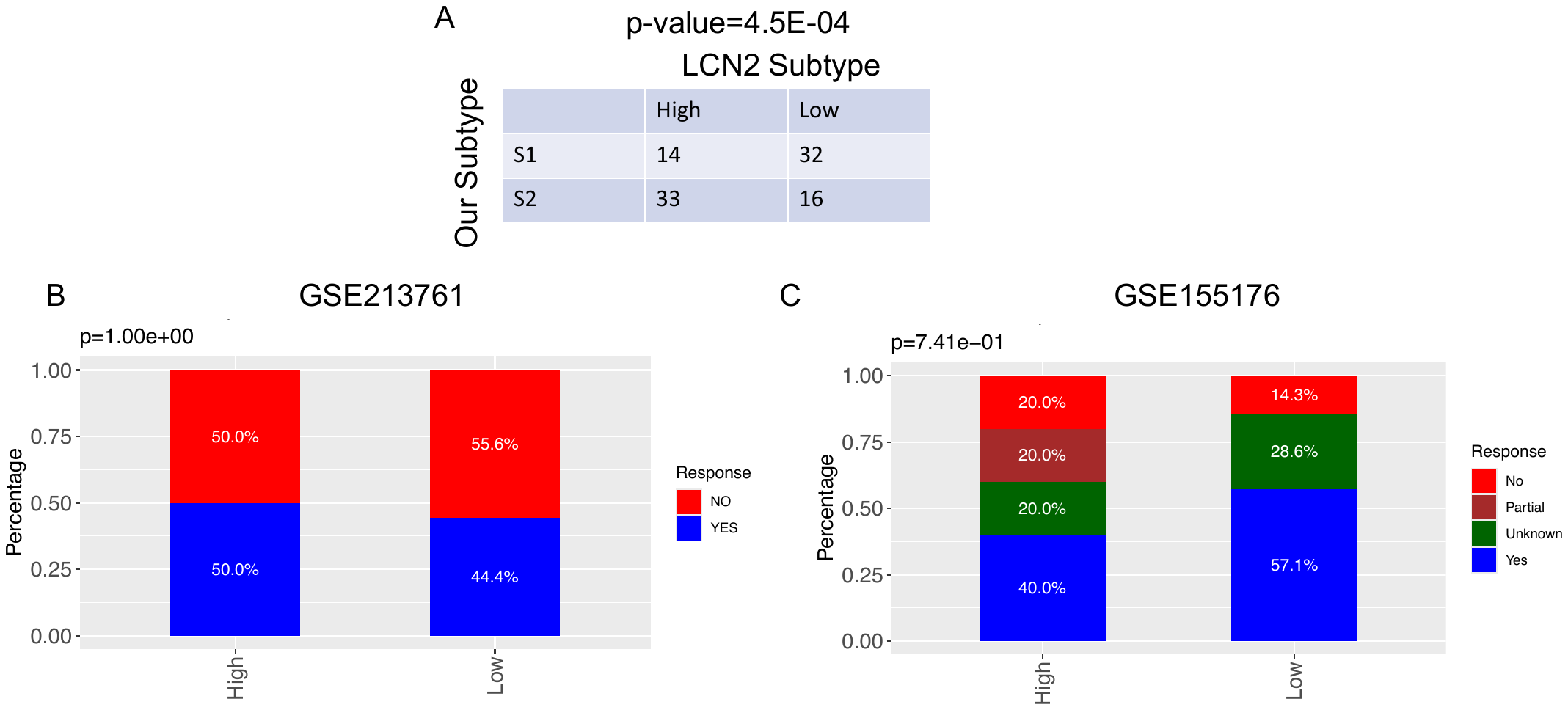

Supplementary Figure S6. Comparison of subtypes identified in this study with LCN2 subtypes (A). (B-C) Relationship between LCN2 subtypes and drug response in GSE213761 and GSE155176. The p-values were calculated by Fisher’s exact test.

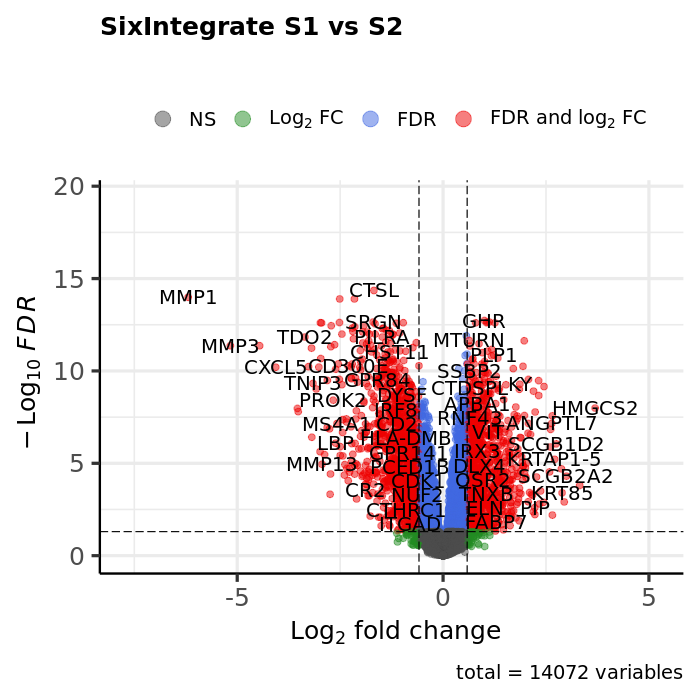

Supplementary Figure S7. Volcano plot for the comparison between subtype S1 and subtype S2. The log2(Fold change) and FDR were calculated by the R limma package.

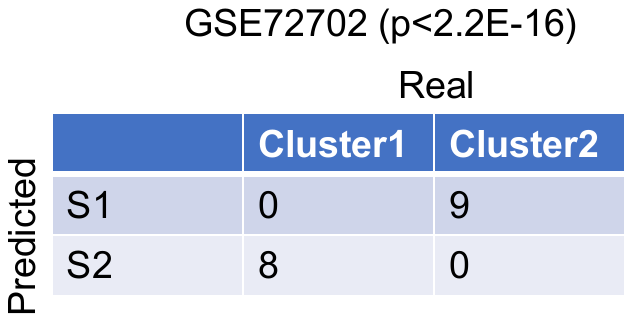

Supplementary Figure S8. Comparison of clusters identified by consensus clustering with subtypes predicted by 36-gene classifier in GSE72702. The p value was calculated by Fisher’s exact test.

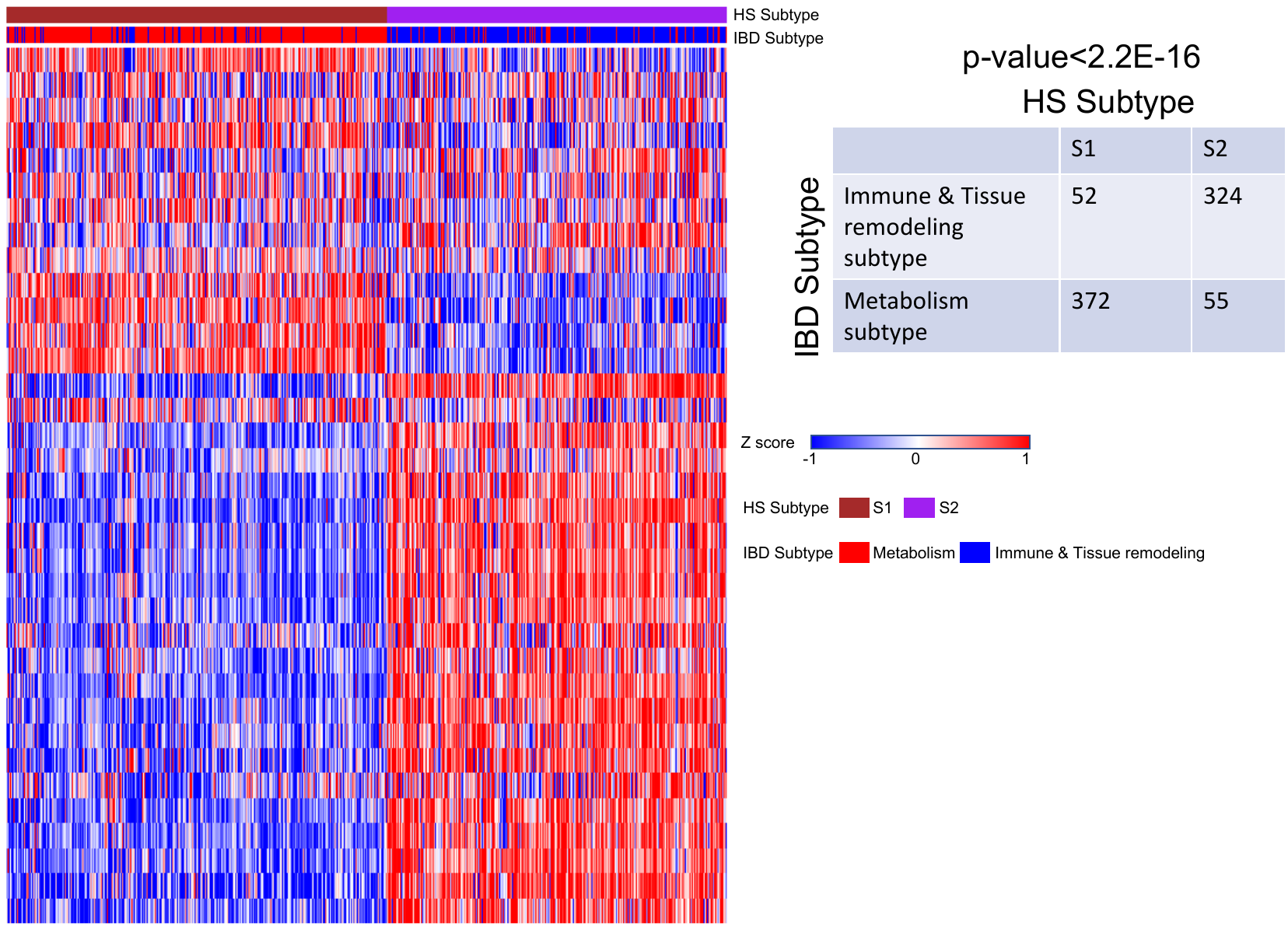

Supplementary Figure S9. Comparison of HS subtypes with IBD subtypes. The p value was calculated by the Fisher’s exact test.
